# Post-translational modifications in the brain are critical contributors to Alzheimer’s disease neuropathology and cognitive decline

**DOI:** 10.64898/2026.06.13.732018

**Authors:** Julia B. Libby, Emily R. Mahoney, Ben Drucker, Philip L. De Jager, Vilas Menon, Shahram Oveisgharan, Julie A. Schneider, Lisa L. Barnes, David A. Bennett, Vladislav A. Petyuk, Timothy J. Hohman

## Abstract

Post-translational modifications (PTMs) in APP and MAPT contribute to plaques and tangles in Alzheimer’s disease (AD). Yet broader proteome-wide PTMs in the AD brain are relatively unexplored. Therefore, this study highlights associations between PTMs, quantified by mass spectrometry in prefrontal cortex tissue, and Alzheimer’s disease neuropathology and cognition.

Leveraging PTMs quantified from prefrontal cortices in 101 Rush Memory and Aging Project participants. We assessed associations with post-mortem amyloid-β and tau burden, global cognition, and cognitive decline. First, APP and MAPT PTM associations were assessed on these outcomes given their known relevance in AD, followed by assessment of protein-wide effects of PTMs. Then, kinase enrichment analysis was performed on each outcome to assess which kinases might contribute to the results.

We observed a novel association of APP-K687 acetylation, a known mutation hotspot driving pathology, with amyloid-β load (β=0.44, P=3.9e-8), while confirming known MAPT PTMs with tangle burden. Further, we identified 20+ novel PTMs’ associations with AD neuropathology, including ENO2-K256 ubiquitination (β=0.353, P=1.13e-6), PSMD13-K31 ubiquitination (β=0.568, P=1.34e-6), and PLXND1-K1826 ubiquitination (β=0.577, P=7.08e-8) for tangle burden and SYP-K23 ubiquitination (β=1.50, P=4.7e-8), TMEFF2-C80 cysteine oxidation (β=1.64, P=1.1e-8), and STX1B-T121 phosphorylation (β=0.898, P=3.3e-7) for amyloid-β load. Further, kinase enrichment analyses highlight the complexity of disease-related proteome changes with some kinases like CDK5 showing expected over-enrichment (amyloid z=3.44, P=3.0e-4; tau z=4.98, P=3.3e-7) but others like PKC family kinases showing divergent enrichment between amyloid (z=8.98–11.55, P<1.0e-18) and tau (z=-2.83–-3.88, P<0.006).

This study provides an atlas of brain PTMs within crucial proteins like MAPT and APP and at the proteome-wide level, that impact AD neuropathology and clinical presentation. Further, we explored what kinases might be driving phosphorylation results, emphasizing the complex proteome changes which impact AD. In sum, these results highlight robust post-translational alterations in the AD brain and provide novel targets for future mechanistic studies.

## Introduction

Following translation, proteins may undergo many changes in structure and function that modify their function and downstream biological effects.(1) Because post-translational modifications (PTMs) play such a pivotal role in protein function, any additional or missing PTMs for a protein can have detrimental effects on a protein’s ability to fold, signal, or bind and can be a key contributor to protein aggregation in disease.(2,3) Indeed, PTMs are known to be associated with protein aggregation in numerous neurodegenerative diseases including Parkinson’s disease (PD), Huntington’s disease (HD), amyotrophic lateral sclerosis (ALS), and Alzheimer’s disease (AD).(2)

AD is characterized by aggregates of amyloid-β and hyperphosphorylated tau tangles, directly impacted by PTMs of the amyloid precursor protein (APP) and microtubule associated protein tau (MAPT), respectively.(4,5) APP is cleaved by either α-secretase (like ADAM10) or β-secretase (BACE1) followed by γ-secretase, and any resulting short amyloid-β peptides are cleared from the brain under healthy conditions.(6,7) However, in disease conditions, this normal processing is disrupted, resulting in an overproduction of amyloid-β in the case of familial AD or a failure to clear amyloid-β in the case of sporadic AD, leading to the accumulation of amyloid-β plaques throughout the cortex.(6–8) Similarly, MAPT is an essential protein in healthy conditions, where tau plays an important role in the assembly, maintenance, and stability of microtubules in neurons.(9) MAPT is phosphorylated at numerous sites throughout the protein, which is required for its interaction with tubulin and its role in microtubule maintenance and assembly.(10) However, in disease states, tau is hyperphosphorylated, leading to aggregation, neurofibrillary tangle development within the neuron, and ultimately neuronal death.(11) Thus, understanding the PTMs in these two critical proteins over the course of AD pathogenesis is critical to uncovering novel pathways for intervention.

The PTMs to APP and MAPT are only beginning to be comprehensively characterized. Multiple PTMs of APP and MAPT have been shown to be associated with their respective protein aggregates like phosphorylation,(12) acetylation,(13) ubiquitiniation,(14) and many more.(15,16) However, most PTM studies of APP and MAPT have been limited to cell and animal models and have characterized the effects of a single modification in detail within these models. Our work will build on this important literature by providing a comprehensive overview of ubiquitination, phosphorylation, acetylation, and cysteine oxidation of these two critical proteins in the context of AD.

Moreover, the proteomic landscape of AD is more complex than APP and MAPT. Many proteins interact directly with APP and MAPT and drive AD neuropathology, such as α-secretase, β-secretase, and γ-secretase as described above, along with other proteins that drive aggregation and hyperphosphorylation.(17) Novel PTMs to proteins that are yet to be characterized could also contribute to protein aggregation or the downstream response to protein aggregation, driving the clinical manifestation of AD. Therefore, the goal of this study is to present proteome-wide characterization of the effects of multiple PTMs including phosphorylation, acetylation, ubiquitination, and cysteine oxidation within brain samples from older adults across the AD spectrum to characterize PTMs that relate to the neuropathological presentation and clinical progression of AD.

## Materials and Methods

### Participants

Data were obtained from the Rush Memory and Aging Project (MAP).(18) MAP is a cohort study of older adults without known dementia who agree to annual clinical evaluations and brain donation at death with recruitment running from September 1997 to the present.(18) The study was approved by the Institutional Review Board (IRB) of Rush University Medical Center. All participants signed informed and repository consents, and an Anatomic Gift Act. Data were downloaded in June 2024. Secondary analyses in this study were approved by the Vanderbilt University Medical Center IRB.

### MS Sample Processing and Data Collection

#### Protein extraction and Lys-C/trypsin tandem digestion

The starting material was a 20-30 mg punch biopsy of human frozen dorsolateral prefrontal cortex (DLPFC) tissue from 103 donors. Samples were homogenized in 100 uL of the lysis buffer (8 M urea, 75 mM NaCl, 50 mM Tris-HCl pH 8.0, 1 mM EDTA, 2 mg/mL Apotinin, 10 mg/mL Leupeptin, 1 mM PMSF, 10 mM NaF, 10 mM sodium butyrate, 50 mM PR-619, and phosphatase inhibitor cocktails 2 and 3 according to the manufacturer’s recommended dilutions) using motorized pestle followed by vortexing at the maximum setting for 10 seconds. Lysates were precleared by centrifugation at 20,000<u>g</u> for 10 min at 4 °C. After the supernatants were transferred to new tubes the protein concentrations were determined by BCA assay. Samples were diluted with the lysis buffer to adjust the protein concentrations to 8 mg/mL. Proteins were reduced with 5 mM dithiothreitol (DTT) for 1 h at 37 °C and subsequently alkylated with 10 mM iodoacetamide for 45 min at 25°C in the dark (both steps were done in ThermoMixer with 1000 rpm shaking). Samples were diluted 1:3 with 50 mM Tris-HCl, pH 8.0 and digested first with Lys-C (Wako) at 1:50 (mAU/ug) enzyme-to-substrate ratio for 2 h at 25 °C, then by sequencing grade modified trypsin (Promega, V5117) at 1:50 enzyme-to-substrate ratio for 14 h at 30 °C. The digested samples were then acidified with 100% formic acid to 1% of the final concentration. Samples were further diluted with 0.1% formic acid to 1.5 mL volume, cleared from the insoluble material by centrifugation for 15 min at 1,500g followed by transfer of the supernatant to new tubes. Tryptic peptides were desalted on C18 SPE (Waters tC18 SepPak, cat. no. WAT054925) using a standard protocol(19) and dried using Speed-Vac. Peptides stored dried at −80 °C till the next step.

#### Pooled reference sample creation

On average the recovered peptide amount was 809 mg. A pooled reference sample was created by taking an equal volume from each of the samples. Further processing of the pooled sample is identical to the subject samples. It was labeled with TMT-16 reagent corresponding to the 126 channel and was used as reference during the FragPipe data analysis.(20) The exception is cysteine oxidation sample analysis that started with independent tissue biopsies (described below).

#### Ubiquityl peptide enrichment

The enrichment was performed using PTMScan HS Ubiquitin/SUMO Remnant Motif (K-ε-GG) Kit (Cell Signaling Technology, cat. no. 59322) according to the manufacturer’s recommendation. Briefly, up to 750 µg of peptide from each sample was reconstituted in 750 µL in PTMScan HS Immunoaffinity Purification (IAP) Bind Buffer. The solution was cleared by centrifugation at 10,000g for 10 min at room temperature. The peptides were then added to 10 µL of 1 x PBS pre-washed PTMScan HS anti-K-ε-GG magnetic bead slurry. The tubes were incubated for 1 h at 4 °C with gentle shaking. The unbound peptide fraction (“flow through”) was collected for further analyses as global proteome and further PTM enrichment. The beads with the bound K-ε-GG peptides were washed with 1 mL of IAP buffer and then with 1 mL of water. After washing, the K-ε-GG peptides were labeled on-bead with 400 mg TMT-16 (ThermoFisher Scientific) in 200 mL of 100 mM HEPES pH 8.5 for 15 min at room temperature with gentle shaking at 1400 rpm. The labeling was quenched by adding 8 mL of 5% hydroxylamine and incubation with shaking for 15 min at room temperature. The beads were washed three times with 1 mL of IAP wash buffer, then eluted by incubating with 50 mL of 0.15% TFA in ice for 10 minutes. The beads were collected at the bottom by centrifugation at 2,500g for 1 minute. The supernatants of the individual TMT-labeled samples were then combined into their respective TMT-16 plexes. Eluates were desalted with C18 StageTips, frozen, and dried in Speed-Vac. For LC-MS/MS analysis, TMT-labeled K-ε-GG peptides were reconstituted in 20 mL of 3% acetonitrile (ACN), 0.1% formic acid (FA), and 0.01% n-Dodecyl β-D-maltoside (DDM). The remaining flow through (non-labeled, K-ε-GG depleted) peptide samples were acidified to a final concentration of 1% FA and desalted on Sep-Pac tC18 SPE, concentrated in Speed-Vac, and the peptide concentration was measured by BCA Assay. The flow through fraction was stored frozen until further processing. The TMT-16 plexes of the K-ε-GG enriched peptides were analyzed by LC-MS/MS “as is”, that is without fractionation.

#### TMT-16 labeling of the global proteome

Samples were labeled with 16-plex TMT reagents using conditions modified from the manufacturer’s instructions (ThermoFisher Scientific). Peptides (400 mg) from each of the samples were dissolved in 80 mL of 50 mM HEPES, pH 8.5 solution. TMT reagents were added at 400 mg amount (20 mL of 20 mg/mL dissolved in anhydrous acetonitrile). After 1 h incubation at 25 °C with 1000 rpm shaking each sample was diluted with 60 mL 100 mM HEPES pH 8.5, 20% acetonitrile. Quench of the reaction was achieved by adding 12 mL of 5% hydroxylamine and incubating for 15 min at 25 °C. Peptides labeled by different TMT reagents were then mixed, dried using Speed-Vac, reconstituted with 3% acetonitrile, 0.1% formic acid and were desalted on tC18 SepPak SPE columns.

#### Multiplexed sample fractionation

Approximately 6 mg of 16-plex TMT labeled sample was separated on a reversed-phase Agilent Zorbax 300 Extend-C18 column (250 mm x 4.6 mm column containing 3.5-mm particles) using Agilent 1200 HPLC System. Solvent A was 5 mM ammonium formate, pH 10, 2% acetonitrile and solvent B was 5 mM ammonium formate, pH 10, 90% acetonitrile. The flow rate was 1 mL/min and the injection volume was 900 mL. The LC gradient started with a linear increase of solvent B to 16% in 6 min, then linearly increased to 40% B in 70 min, 4 min to 44% B, 5 min to 60% B and another 14 of 60% solvent B. A total of 96 fractions were collected into a 96 well plate throughout the LC gradient. These fractions were concatenated into 24 fractions by combining 4 fractions that are 24 fractions apart (i.e., combining fractions #1, #25, #49, and #73; #2, #26, #50, and #74; and so on). For global proteome analysis, 5% of each concatenated fraction was dried down and re-suspended in 2% acetonitrile, 0.1% formic acid to a peptide concentration of 0.1 mg/mL for LC-MS/MS analysis. The rest of the fractions (95%) were further concatenated into 12 fractions (i.e., by combining fractions #1 and #13; #3 and #15; and so on), dried down, and subjected to immobilized metal affinity chromatography (IMAC) for phosphopeptide enrichment followed by lysine acetylation enrichment.

#### Phosphopeptide enrichment using IMAC

Fe3+-NTA-agarose beads were freshly prepared using the Ni-NTA Superflow agarose beads (QIAGEN, #30410) according to the manufacturers recommended protocol. For each of the 12 fractions, peptides were reconstituted to ∼0.5 mg/mL in 500 mL IMAC binding/wash buffer (80% acetonitrile, 0.1% trifluoroacetic acid) and incubated with 40 mL of the 50% bead suspension for 30 min at 25 °C. After incubation, the beads were washed 2 times each with 50 mL of wash buffer and once with 50 mL of 1% formic acid on the stage tip packed with 2 discs of Empore C18 material (Empore Octadecyl C18, 47 mm; Supleco, 66883-U). Phosphopeptides were eluted from the beads on C18 using 70 mL of Elution Buffer (500 mM K_2_HPO_4_, pH 7.0). 50% acetonitrile, 0.1% formic acid was used for elution of phosphopeptides from the C18 stage tips. Samples were dried using Speed-Vac and later reconstituted with 10 mL of 3% acetonitrile, 0.1% formic acid for LC-MS/MS analysis.

#### Acetyl-lysine immunoaffinity enrichment

Phosphopeptide-depleted flowthroughs were concatenated into 6 fractions by combining 2 fractions that are 6 fractions apart (i.e., combining fractions #1 and #7 as a new fraction), dried using a Speed-Vac, and reconstituted in 250 μL of IAP Bind Buffer (Cell Signaling Technology) containing 0.01% CHAPS. Antibody magnetic beads from the PTMScan Acetyl-Lysine Motif [Ac-K] Kit (Cell Signaling Technology, cat. no. 13416) were freshly prepared for each enrichment by centrifugation, removal of storage buffer, and washing four times with 1 mL of ice-cold 1× PBS, followed by resuspension in IAP Bind Buffer with 0.01% CHAPS. Five microliters of bead slurry were added to the wells of the KingFisher plate, peptides were transferred to the corresponding wells, and the plate was incubated for 3 h at 4 °C on a rotator. Bead-bound acetyl peptides were washed four times with 250 mL HS IAP Wash Buffer containing 0.01% CHAPS, followed by one wash with 0.01% CHAPS solution. Bound peptides were eluted with 100 mL of 0.15% TFA after 10 min incubation at room temperature. The enriched acetyl-lysine peptides were purified by desalting on C18 StageTips. Followed by elution with 50% acetonitrile/0.1% formic acid. The eluates were dried in a Speed-Vac and reconstituted in 12 mL of 3% acetonitrile, 0.1% formic acid, with 0.01% n-Dodecyl β-D-maltoside (DDM), prior to LC-MS/MS analysis.

#### Enrichment of the cysteine oxidation

Dorsolateral prefrontal cortex brain biopsy samples (∼30 mg) were processed as previously described. (21,22) Samples were processed one TMT plex (14-15 samples) at a time. Samples were quickly minced on a glass slide on dry ice and placed in 5 mL round bottom tube. Small portions of tissue from each of the samples were combined to create an additional “total thiol” sample for each plex. A volume of 700 mL of homogenization buffer (250 mM MES pH 6.0, 1% SDS, 1% Triton-X, 5 mM EDTA) was added to each of the samples. Buffer for all samples except “total thiol” included 100 mM N-ethylmaleimide (NEM) to block free (reduced) thiol groups. To allow the NEM to block thiols, the samples were incubated for 30 min at room temperature in the dark prior to the homogenization. The step was followed by homogenization with a Tissue-Tearor probe homogenizer (BioSpec Products, Bartlesville, OK). Homogenates were transferred to 1.5 mL tube, centrifuged at 16,000g at 4 °C for 10 min to remove insoluble material. Supernatant was transferred to 5 mL safe-lok tubes. For the denaturation step, samples were incubated in the dark at 55 °C for 30 min with 850 rpm shaking. Proteins were precipitated by adding 4x volume of acetone at −20 °C, vortexing, and incubating overnight at −20 °C. Protein pellets were collected by centrifugation for 10 min at 13,000 rpm. The pellets were washed twice with 3 mL ice cold acetone. The protein pellets were resuspended in denaturation buffer (50 mM Tris-HCl pH 8.0, 8 M urea). Protein concentrations were measured using BCA assay and 400 mg aliquots were taken for further processing. Samples were supplemented with 10 mM DTT and incubated for 1 hour at 37 °C with 1000 rpm shaking. Samples were diluted 4x with 50 mM Tris-HCl and digested with sequence-grade trypsin (Promega) at a 1:50 (mg trypsin: mg protein) ratio for 1 h at 37 °C, followed by a 2 −fold dilution with 50 mM Tris-HCl and additional trypsin (to maintain 1:50 ratio) and digested overnight. The peptides were cleaned up by C-18 SPE and 100 mg of peptide from each sample was labeled with TMT-18 reagents (ThermoFisher). 134N, 134C and in some plexes 133C were not used for quantification of total oxidation due to potential interference from the 135N tags used to quantify total thiol. The labeled peptides were then resuspended in 250 mM HEPES pH 8, reduced with DTT at a final concentration of 5 mM, and a 100 mg aliquot of peptide was collected for global proteomics. The remaining peptide was diluted 6-fold with 25 mM HEPES pH 7.7 to obtain a DTT concentration of ∼0.8 mM to permit enrichment of Cys-containing peptides with thiol affinity resin. The enrichment procedure is adapted from our RAC-TMT workflow.(23) After enrichment, the enriched “oxidized thiol” peptides and global peptides were alkylated with iodoacetamide at 4 times the molarity of the DTT present in the samples and desalted using C-18 SPE. Aliquots of 20 mg of global or enriched redox peptides were collected into 12 concatenated fractions as previously described.(21) Fractions were submitted for LC-MS/MS analysis at 0.1 mg/mL concentration in 0.1% formic acid.

#### LC-MS/MS data acquisition

Fractionated samples prepared for global proteome, phosphoproteome, acetylome, and cysteine oxidation analysis were separated using a nanoACQUITY UPLC system (Waters) by reversed-phase HPLC. The analytical column was manufactured inhouse using ReproSil-Pur 120 C18-AQ 1.9 mm stationary phase (Dr. Maisch GmbH) and slurry packed into a 25-cm length of 360 mm o.d. x 75 mm i.d. fused silica picofrit capillary tubing (New Objective). The analytical column was heated to 50 °C using an AgileSLEEVE column heater (Analytical Sales and Services). The analytical column was equilibrated to 98% mobile phase A (MPA, 0.1% formic acid/3% acetonitrile) and 2% mobile phase B (MPB, 0.1% formic acid/90% acetonitrile) and maintained at a constant column flow of 200 nL/min. The sample was injected into a 5-mL loop placed in-line with the analytical column which initiated the gradient profile (min:%MP B): 0:2, 1:6, 85:30, 94:60, 95:90, 100:90, 101:50, 110:50. The column was allowed to equilibrate at start conditions for 30 minutes between analytical runs.

Analysis was performed using an Orbitrap Fusion Lumos mass spectrometer (ThermoFisher Scientific). Electrospray voltage (1.8 kV) was applied at a carbon composite union (Valco Instruments) coupling a 360 mm o.d. x 20 mm i.d. fused silica extension from the LC gradient pump to the analytical column and the ion transfer tube was set at 250 °C. Following a 25 min delay from the time of sample injection, Orbitrap precursor spectra (AGC 4 x 10^5^) were collected from 350-1800 m/z for 110 min at a resolution of 60K along with data dependent Orbitrap HCD MS/MS spectra (centroid) at a resolution of 50K (AGC 1 x 10^5^) and max ion time of 105 ms for a total duty cycle of 2 seconds. Masses selected for MS/MS were isolated (quadrupole) at a width of 0.7 m/z and fragmented using a collision energy of 30%. Peptide mode was selected for monoisotopic precursor scan and charge state screening was enabled to reject unassigned 1+, 7+, 8+, and > 8+ ions with a dynamic exclusion time of 45 seconds to discriminate against previously analyzed ions between ±10 ppm.

#### LC-MS/MS data analysis

The global and phosphoproteomics datasets were analyzed with FragPipe tool(20) with essentially default settings: TMT16, TMT16-phospho, TMT16-acetyl or TMT16-Ubi workflows. The ubiquitination workflow additionally allowed the presence of TMT label on the K-GG residue and required minimum 0.5 site localization probability for quantification. The protein sequence database was represented by canonical sequences UniProt/SwissProt v 2021-06-20 with exception of the gene MAPT. In the case of MAPT, the canonical form P10636, which isn’t expressed in the brain, was replaced with the P10636-8 (2N4R) isoform, which is conventionally used as a reference isoform for tau phosphorylation studies. Global and PTM data were summarized at gene and single site levels, respectively. All the data was reported as log2-transformed abundances relative to the reference sample. The normalization by FragPipe was turned off and was performed later using preprocessing R scripts. The normalization coefficients accounting for the sample-to-sample amount differences for the global samples were computed based on the assumption that the proteome abundance overall shouldn’t change. Then, those normalization coefficients derived from the global proteome were applied to the PTM samples. The exception was ubiquitination, since the TMT labeling was done independently of the global proteome. Batch effects were corrected both in global and phospho datasets using the ComBat approach.(24) To isolate the effect of the stoichiometry of the PTMs, the change of the parent proteins was regressed out of the PTM relative abundances.

### Cognitive Measures

Cognitive performance was annually quantified using 19 cognitive tests annually spanning five cognitive domains including working memory, episodic memory, semantic memory, percentual speed, and visuospatial ability. A global cognition composite was calculated using the average of z-scores from the 19 tests. Further methods have been described elsewhere.^23^ A global cognition slope was calculated as the estimated person-specific rate of change in the global cognition variable over time. The global cognition slope was calculated by adjusting for demographics including age of death, sex, and years of education with time modeled as years from baseline. An additional slope was calculated adjusting for multiple pathologies in addition to demographics (age of death, sex, years of education) with time modeled as years from death.(26)

### Measures of Alzheimer’s Disease Pathology

MAP pathology measures at autopsy have been described elsewhere.(18) To summarize, tau tangles and amyloid-β deposits were immunohistochemically quantified across 8 regions including the hippocampus, angular gyrus, and entorhinal, midfrontal, inferior temporal, calcarine, superior frontal, and anterior cingulate cortices measured. Regional measures were scaled and averaged to yield tau tangle density and amyloid-β load. Final measurements were square-root transformed to approximate a normal distribution.

### Statistical Analyses

Statistical analyses were completed using R (version 4.3.0) and code is available from the authors upon request. P-values were corrected for multiple comparisons using the false discovery rate (FDR) procedure and significance was set *a priori* at α=0.05 FDR.(27)

Of the 103 participants with PTM data, two were dropped who had non-AD dementia leaving 101 for analysis. Linear regressions (covarying for sex, age at death, APOE-ε4 carrier status, and post-mortem interval) tested PTM protein site associations with cross-sectional global cognition, slopes of global cognition adjusted for demographic variables, tau tangle density, amyloid-β load, and clinical diagnosis at death. For longitudinal analyses, the slope outcome is preadjusted for the effects of age at baseline, sex, and education so these variables were not included as additional covariates in those analyses.

Analyses first evaluated APP and MAPT sites given their known importance in AD pathogenesis. Then, a proteome-wide discovery analysis looked for novel PTM site associations with AD outcomes. Within APP and MAPT specific analyses, FDR corrections were carried out across all PTMs and outcomes. For proteome-wide analyses, FDR corrections were carried out across all PTMs within each outcome.

Additionally, phosphorylation kinase enrichment analysis (KSEA)(28–30) was run for each outcome using the “KSEA App” implementation for R.(31) We aimed to obtain a large set of kinase-phosphorylation site pairings that was broader than the default PhosphositePlus(29) database of approximately 22k kinase-site pairings. To this end, we used the Kinase Library Python tool,(32) a program which predicts phosphorylation events between human kinases and arbitrary amino acid substrate sequences. As input to KSEA, we included putative kinase-site pairings with a “site percentile” prediction score (as defined by Kinase Library) above 90%. Ultimately, our KSEA input consisted of over 600k kinase-site pairings. This allowed us to determine, comprehensively, which kinases contributed to the observed results. For interpretability, the direction of cognitive outcomes was inverted so that higher indicated worse cognition or faster decline while lower indicated better cognition and slower decline. FDR correction for p-values was applied to KSEA results within each outcome with a threshold of 5%.

## Results

Data from the Rush Memory and Aging Project (ROS/MAP)(18) were leveraged for this study, including annual cognitive measures, post-mortem quantification of tau tangle density and amyloid-β load, as well as four types of PTMs (ubiquitination, phosphorylation, acetylation, and cysteine oxidation). See the **Online Methods** for more details. Participant characteristics are presented in **Table 1**. The 101 participants were old, highly educated individuals with the majority cognitively impaired and female.

**Table 1.** Descriptive Statistics.

|  | NCI<br>(N=34) | MCI<br>(N=33) | AD<br>(N=34) | Total<br>(N=101) | P-value |
| --- | --- | --- | --- | --- | --- |
| Male, no. (%) | 12 (35) | 12 (36) | 14 (41) | 38 (38) | 0.027 |
| APOE-E4 Carriers, no. (%) | 3 (9) | 10 (30) | 12 (35) | 25 (25) | 0.868 |
| Age of death, yrs | 88.2 +/- 6.3 | 88.9 +/- 5.1 | 91.0 +/- 4.8 | 89.4 +/- 5.5 | 0.108 |
| Education, yrs | 14.5 +/- 2.7 | 14.8 +/- 2.2 | 15.4 +/- 2.9 | 14.9 +/- 2.6 | 0.431 |
| Post-mortem interval, hrs | 5.7 +/- 1.8 | 7.2 +/- 4.1 | 6.8 +/- 3.1 | 6.6 +/- 3.2 | 0.129 |
| Global cognition composite (at last visit), z | 0.1 +/- 0.5 | -0.6 +/- 0.4 | -1.9 +/- 0.9 | -0.8 +/- 1.0 | < 0.001 |
| β-amyloid load | 1.2 +/- 1.1 | 1.6 +/- 1.3 | 2.5 +/- 1.1 | 1.8 +/- 1.3 | < 0.001 |
| Tau tangle density | 1.4 +/- 0.7 | 2.1 +/- 1.2 | 3.0 +/- 1.3 | 2.2 +/- 1.3 | < 0.001 |

### APP and MAPT PTMs

First, given their known importance in AD pathogenesis, associations of APP and MAPT PTM sites were assessed with cross-sectional global cognition, slopes of global cognition adjusted for demographic variables, slopes of global cognition adjusted for demographic variables and post-mortem pathologies, tau tangle density, amyloid-β load, and clinical diagnosis at death. Cross-sectional models were adjusted for sex, age at death, APOE-ε4 carrier status, and post-mortem interval. Longitudinal analyses used slopes preadjusted for the effects of age at baseline, sex, and education so these variables were not included as additional covariates in those analyses. P values were corrected using the false discovery rate (FDR) procedure across all PTMs and outcomes. A summary of all statistically significant results for APP and MAPT passing FDR corrections are presented in **Supplemental Table 1**.

For APP protein sites, acetylation of the K687 site contributed to both amyloid-β load (**Figure 1**) and tangle density. When correcting for amyloid-β peptide abundance rather than just parent protein abundance, the associations between APP-K687 were attenuated (p>0.05). Additionally, cysteine oxidation of site C117 had a higher occurrence in cognitively normal participants.

**Figure 1.**
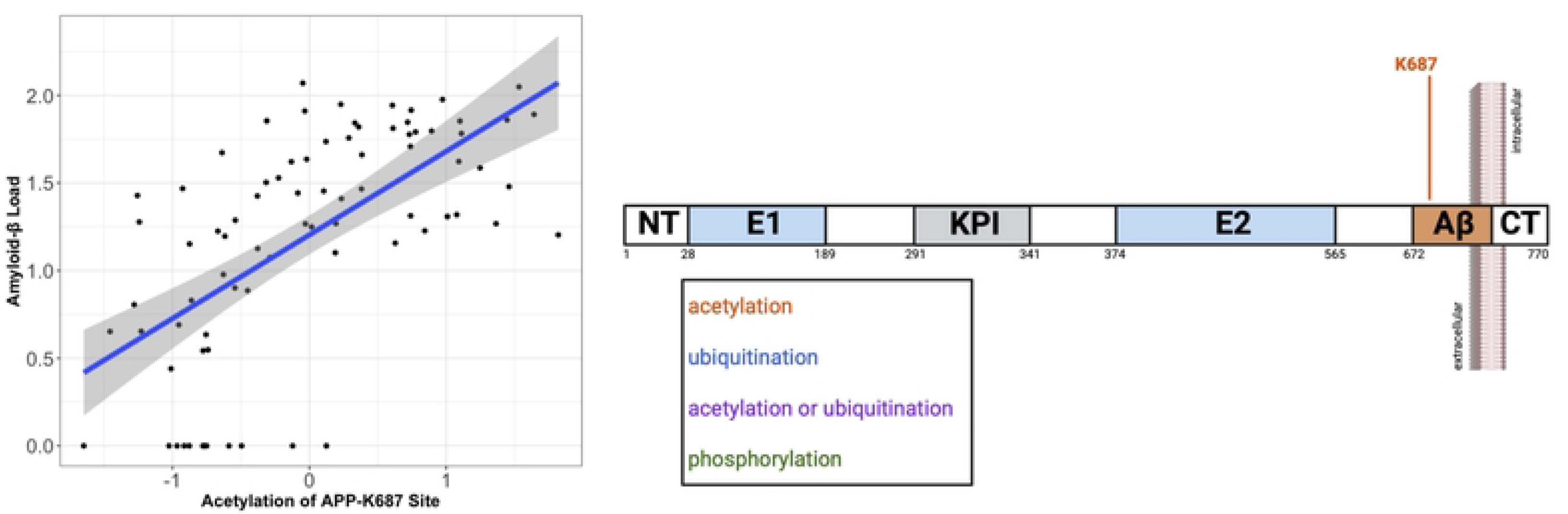
APP-K687 Association with Amyloid-β Load. This figure illustrates the FDR significant signal PTM on APP observed when looking at APP and MAPT sites. (A) Acetylation of the K687 site was associated with a higher amyloid-β load. (B) This site is located within the amyloid-β region of the protein.

For MAPT, 36 PTMs were associated with more tau tangles, illustrated in **Figure 2**, specifically acetylation, ubiquitination, and phosphorylation. Additionally, 13 PTMs were associated with a greater amyloid-β load, 34 with worse cognitive performance at last visit, 31 with a higher rate of cognitive decline over time when controlling for just demographics, and three associated with cognitive impairment. Many of these PTMs had detrimental effects across all outcomes, indicating any modifications to these sites have robust impacts on disease progression. However, interestingly, higher phosphorylation of site T102 was seen among cognitively normal participants compared to AD cases and was associated with better cognitive performance.

**Figure 2.**
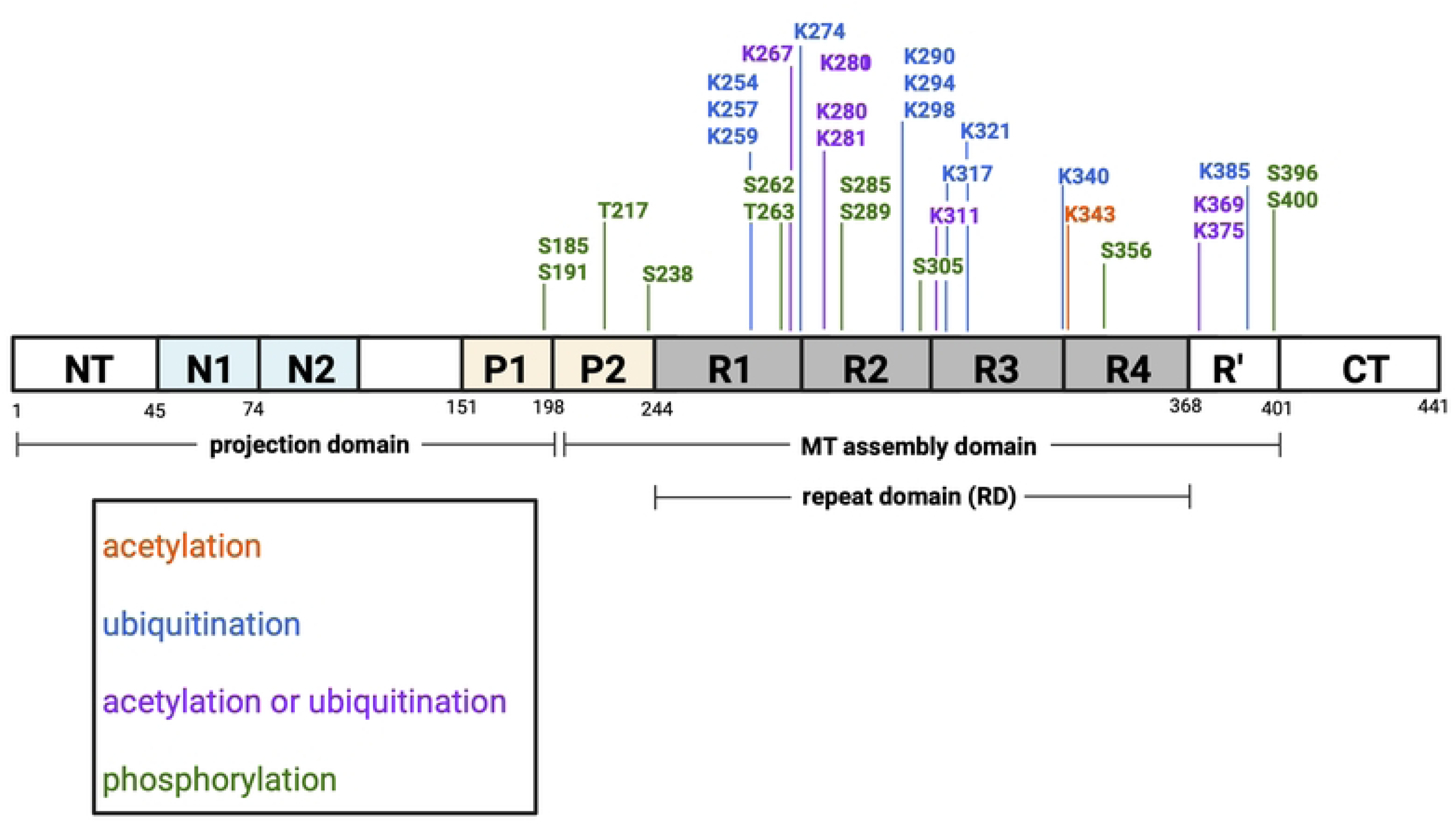
PTMs of MAPT Contributing to Tangle Density. This illustration depicts the sites on the MAPT protein where PTMs were associated with higher tangle density. Each site is colored based on the PTM affecting that site. The numbers underneath the diagram indicate amino acid numbers across the protein.

### Proteome-Wide Effects

Then, proteome-wide associations were assessed in the same manner as the APP and MAPT models. FDR correction was carried out across all PTMs within each outcome. PTMs that passed correction for multiple comparison are stated below. All effect sizes, uncorrected p-values, and corrected p-values are included in **Supplemental Table 2**. Proteome-wide associations that passed FDR correction are illustrated in **Figures 3** and **4**. Many of the APP and MAPT PTM signals remained statistically significant at the proteome-wide level, including the detrimental association of higher acetylation of APP at site K687 on amyloid-β load, and multiple PTMs of MAPT having detrimental associations with tangle density.

**Figure 3.**
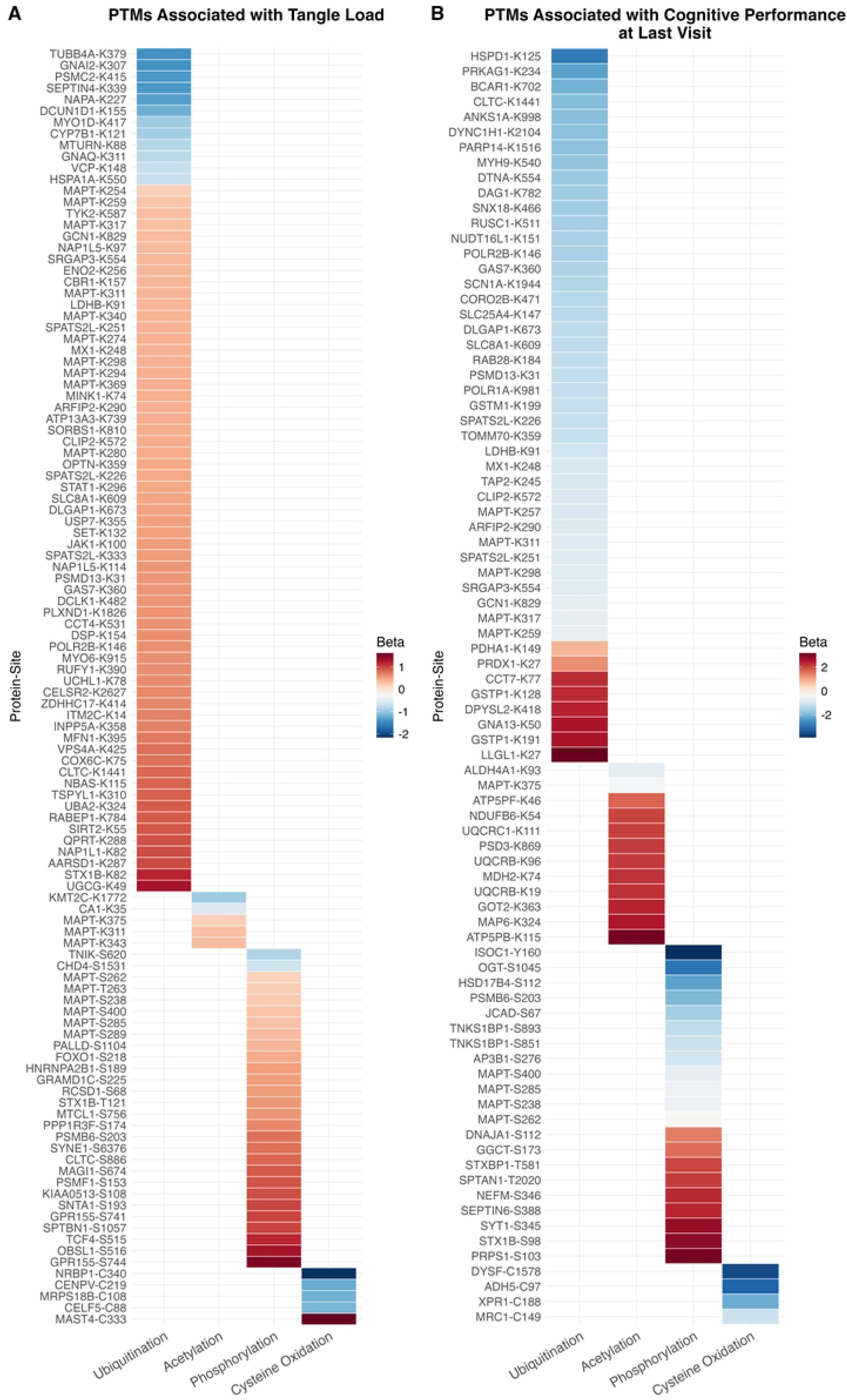
Proteome-Wide PTM Effects on Tangle density and Performance at Last Cognitive Visit. This figure illustrates the proteome-wide associations that pass FDR correction for multiple comparisons for the outcomes (A) tangle density and (B) global cognitive performance at last visit. The x-axis identifies the PTM while the y-axis identifies the specific protein-site where the modification has a significant effect on the outcome.

**Figure 4.**
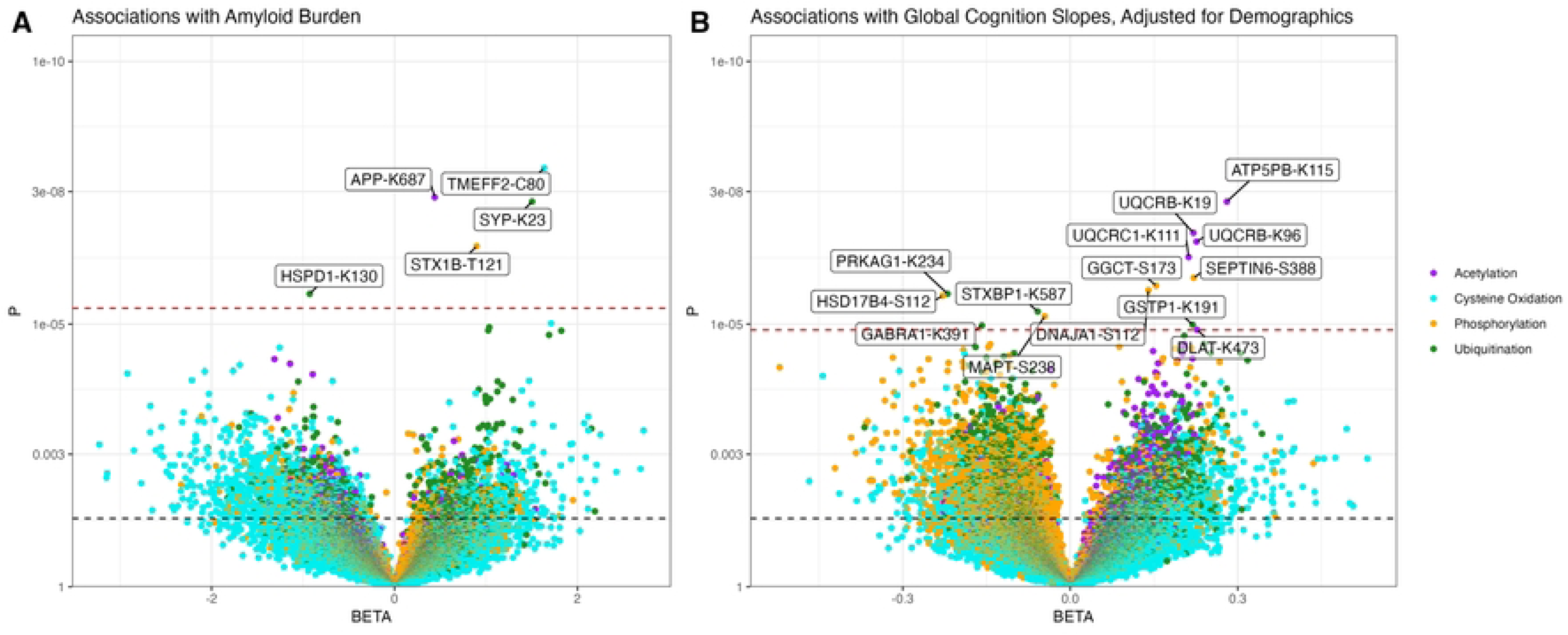
Proteome-Wide PTM Effects on Amyloid-β and Rate of Global Cognitive Decline. This figure illustrates the proteome-wide associations for the outcomes (A) amyloid-β load and (B) the rate of global cognitive decline where the slopes were adjusted for demographic variables. The x-axis represents the effect size of each model, where the y-axis represents the nominal p-value. The black dotted line represents the nominal p-value threshold of 0.05, and the red dotted line represents FDR corrected p-value threshold. Points are colored on PTM.

Higher levels of PTMs across the proteome appeared to relate to more AD neuropathology and worse cognitive function. Robust changes across the proteome were associated with higher tangle density including 92 PTMs (**Figure 3A**), most of which were ubiquitination or phosphorylation. The abundance of three PTMs were associated with higher amyloid-β load in addition to the previously observed APP-K687 signal (**Figure 4A**), including ubiquitination of SYP-K23, cysteine oxidation of TMEFF2-C80, and phosphorylation of STX1B-T121. Additionally, 57 PTMs were associated with worse global cognitive performance at last visit (**Figure 3B**), and 9 associated with a more rapid rate of cognitive decline where cognitive slopes were adjusted for demographic variables (**Figure 4B**). Interestingly, no associations with an Alzheimer’s disease dementia diagnosis survived correction for multiple testing, likely due to the increased statistical power for the quantitative outcomes.

There were some notable protective associations. A higher abundance of 20 PTMs related to a lower tangle density, 27 PTMs were associated with better cognitive performance at the last visit, and 5 PTMs were associated with a slower rate of cognitive decline over time where slopes were adjusted for demographic variables. Additionally, higher ubiquitination of HSPD1 at site K130 was associated with a lower amyloid-β load.

### Kinase Enrichment Results of the Phosphoproteomics Data

Finally, in order to assess which kinases may be the driving forces behind the phosphoproteomic changes in the selected neuropathology and cognition phenotypes, phosphorylation kinase enrichment analyses (KSEA)(28,30,31) was run using Kinase Library(32) kinase-site pairings with a prediction score >90%. KSEA p-values were corrected using the FDR procedure within each outcome. Enriched kinases for each outcome are presented in **Supplemental Table 3**. Interestingly, KSEA demonstrated decreased activity for multiple kinases in relation to tau tangle burden, cognitive performance, and AD dementia, but an over-enrichment for phosphorylation sites related to amyloid-β load (see **Figure 5**). Despite the preponderance of flipped directionality in the kinase activity changes between amyloid-β and the rest of the outcomes, one kinase (VRK2) showed under-enrichment for all outcomes. Further, some kinases showed a consistent direction between amyloid and tau including six kinases that were down-regulated (ACVR1B, ACVR2A, ACVR2B, ACVRL1, CSNK1D, and GRK3) and nine of which were up-regulated, including DYRK1A, PINK1, MOK, HIPK3, MTOR, CDK9, CDK3, CDKL5, and MAPK15. Kinases that are known to interact with tau were enriched for more positive associations with tau, including several kinases in the NEK, PRK, and MAPK families. Moreover, 130 out of 232 of the significantly enriched kinases for tau were driven in part by MAPT PTMs. Numerous novel kinases were also enriched for tangle burden including NLK, UHMK1, and IRAK4.

**Figure 5.**
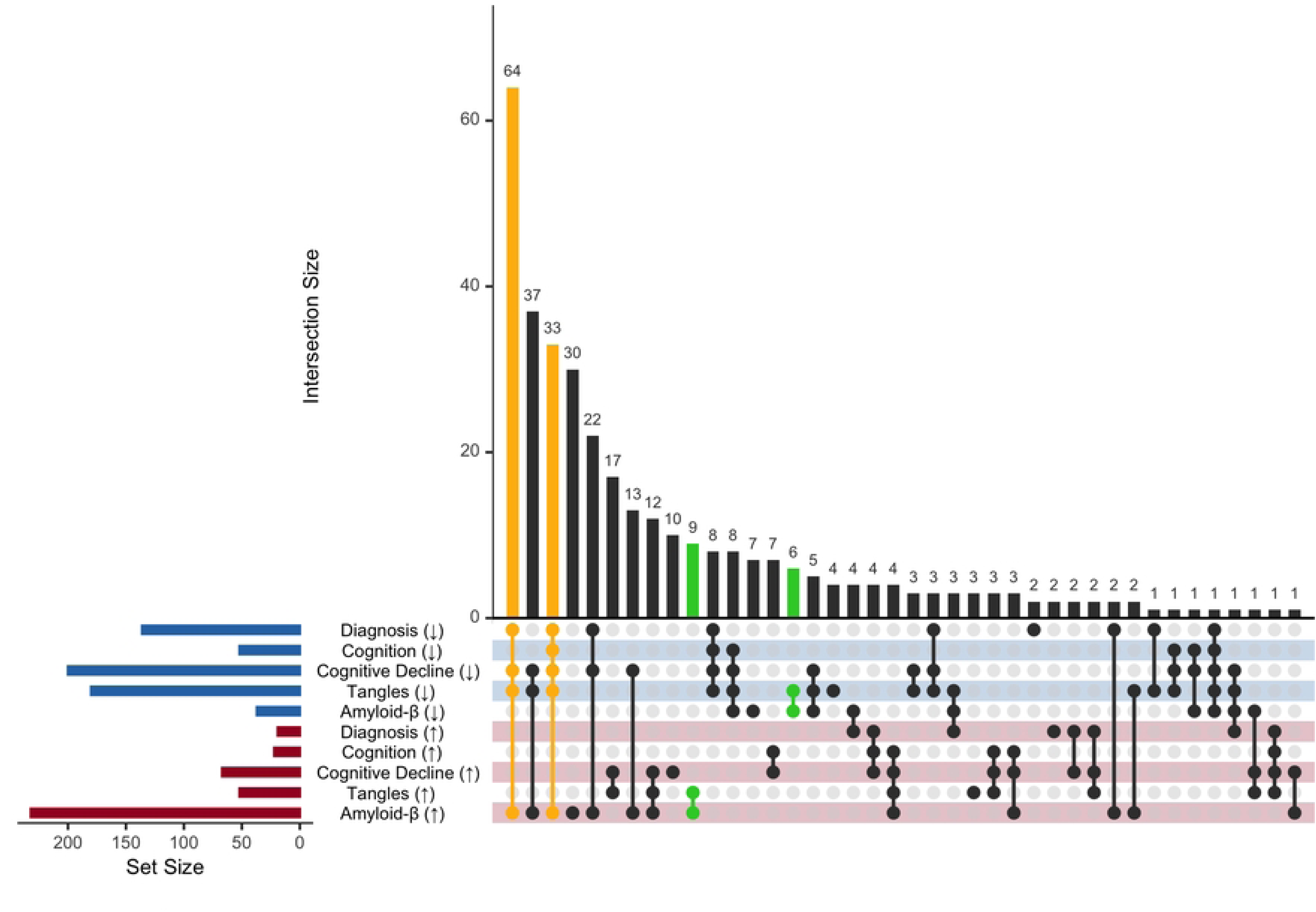
Kinase Enrichment Results Across Outcomes. This figure illustrates the distribution of over-and under-enriched kinases across outcomes. Blue rows indicate down-regulated kinases while red rows indicate up-regulated kinases. Overlap columns highlighted in yellow are shared across all outcomes – cognitive (either cross-sectional or longitudinal), diagnosis, amyloid-β load, and tau tangle density – but with flipped direction in amyloid. Overlap columns highlighted in green are enriched in a consistent direction between amyloid-β and tau tangles. Note that, prior to enrichment analyses, effect directions were inverted on cognitive outcomes for interpretability.

## Discussion

In this study, we analyzed proteome-wide effects of multiple PTMs in the human brain on AD pathology and cognitive decline. We successfully replicated numerous known PTMs in the MAPT protein that relate to tau tangle density, while also uncovering multiple novel PTMs in both APP and MAPT in relation to plaques and tangles, respectively, including acetylation of APP at K687, a known mutation site that causes familial AD.(33–36) We also provide compelling evidence of widespread effects of PTMs on AD outcomes, with a particularly striking number of associations with tangle density. Finally, there were numerous kinases that showed an enrichment for associations with amyloid-β, but an under enrichment for associations with tau tangle burden, suggesting key differences in the way the same kinases contribute to the two neuropathological hallmarks of AD.

We observed novel evidence that acetylation of APP at K687 is related to elevated amyloid-β load. This particular site has not been previously implicated in PTM analysis, but has substantial evidence as a site of detrimental mutations that drive AD.(35–38) Due to K687 being a cleavage site in the non-amyloidogenic APP pathway, mutations at this site can prevent α-secretase from binding properly, disrupting healthy processing of APP.(38) Other studies have shown acetylation of APP on sites like K16,(39) K132 and K134(40) to be associated with lower levels of amyloid-β, believed to be due to acetylation decreasing the binding at those sites and preventing aggregation.(39,40) It is possible that acetylation at K687 similarly decreases α-secretase’s binding affinity, and disrupts non-amyloidogenic processing of APP driving greater amyloid-β aggregation. The observed association may be a consequence of the strong association of this acetylation with the aggregate form, like phosphorylation of α-synuclein at S129.(41) However, like α-synuclein S129 phosphorylation, when this acetylation happens relative to cleavage by the secretases and aggregation is unclear. Further exploration of K687 and its relation to amyloid-β deposition are needed to parse out whether this could be a potential intervention target. Either way, these results build on the literature implicating K687 described above, while highlighting the need for additional mechanistic studies of this acetylation site in AD.

We also observed numerous novel and established PTMs in MAPT in relation to tangle burden. Hyperphosphorylation is known to drive tangle accumulation, so it is not surprising that we observed strong associations between phosphorylation at numerous sites including S185, S238, S262, T263, S285, S289, S305, and S400 with tangle pathology, all of which are well-established sites of phosphorylation in AD.(42) Additionally, in kinase enrichment analyses, several known tau kinases like several CDKs and MAPKs were enriched for associations with higher tangle burden. In addition to the observed phosphorylation effects, we observed robust effects of acetylation and ubiquitination on MAPT in relation to tangle burden. Other studies have also observed vast effects of these PTMs on MAPT in AD and other tauopathies,(14) with many MAPT sites impacted by either modification.(13,14,42,43) While hyperphosphorylation plays a role in tau aggregation due to the protein conformation changes, it appears acetylation and ubiquitination are also common in the presence of neurofibrillary tangles, though whether these modifications are upstream or downstream of tangle formation is less clear.(13,43) There has been evidence that both acetylation and ubiquitination of tau help mediate tau signaling in healthy conditions, but it is speculated that phosphorylated tau is more susceptible to ubiquitin and acetyl modifications, which could in turn accelerate tau aggregation,(43) or perhaps in healthy conditions lead to degradation of a dysfunctional protein. Our results build on this literature and suggest that better characterization of the functional impact of acetylation and ubiquitination is critical in characterizing how this protein changes in disease.

When expanding the scope of the analysis to the entire proteome, widespread effects on AD neuropathology and cognitive performance were observed. Notably, those PTMs associated with cognitive decline were non-significant after correcting for the effect of pathology, suggesting that those associations are driven by the pathology effects. While some of these proteins have been previously implicated in AD, little work has been done to better understand the mechanistic role of these proteins in pathogenesis. Interestingly, we saw multiple signals from proteins involved in cell signaling and neuronal differentiation on AD neuropathology, including that more ubiquitination of PLXND1 at site K1826 was associated with a higher tangle density. This protein is a plexin receptor for semaphorin molecules, involved in cell signaling and migration during vascular development, most prominently in the heart and central nervous system (CNS).(44) Similar effects were observed for ubiquitination of PSMD13 at site K31, a proteasome regulatory subunit with evidence supporting its role in interacting with microRNA to encourage neuronal differentiation.(45) Disruption of this protein can lead to decreased neuronal development(45) and dysregulated proteostasis in AD pathophysiology.(46) Additionally, PSMD13 is believed to interact with phosphorylated tau in the AD brain.(17)

To build on this observation, our results indicate that ubiquitination is elevated in the face of a higher neurofibrillary tangle burden, particularly within pathways involved in neuronal and vascular development in the CNS. Ubiquitination plays a critical role in proteasomal degradation, autophagy, cell death, DNA damage response, and cell signaling,(47) so it is quite likely that this upregulation reflects key shifts in the proteasome in response to AD pathology. Certainly, within diseases characterized by protein aggregation, increased ubiquitin signaling occurs to drive degradation of these aggregates.(48) However, the proteasome is inefficient at degrading aggregates compared to soluble proteins.(48) Our results suggest that ubiquitination of proteins involved in cell signaling like PLXND1 and protein subunits of the proteasome including PSMD13 may be further impairing their ability to respond to tau aggregates, and thus contribute to a feed-forward loop driving neurofibrillary tangle development. Future mechanistic work should explore the role of ubiquitination of these key proteins in the context of tau tangle formation.

While fewer proteins were implicated in amyloid-β aggregation beyond APP, modifications of some proteins appeared to impede their role in amyloid-β clearance or enhance their ability to disrupt this process. For example, cysteine oxidation of TMEFF2 at site C80 was associated with a greater amyloid-β load. Tomoregulin (TMEFF2) is a brain-enriched transmembrane protein that has been shown to bind APP and enhance its cleavage by α-secretase, further enabling non-amyloidogenic APP clearance.(49) While there is little literature on the effects modifications of TMEFF2 have on amyloid-β clearance, we can speculate cysteine oxidation at site C80 modifies the protein structure or function in a way that reduces its binding affinity to APP, perhaps disrupting its role in enhancing amyloid-β clearance, leading to greater amyloid-β aggregation downstream. Additionally, more ubiquitination of SYP at site K23 was associated with a higher amyloid-β load. Synaptophysin (SYP) is a synaptic vesicle protein that has been reported as downregulated in AD.(50) Interestingly, evidence has shown Aβ42 binds to SYP in AD, and in turn reduces neurotransmitter release and synaptic plasticity.(51) One testable hypothesis from our results is that ubiquitination at site K23 may increase SYP’s binding affinity with Aβ42, and thus increase amyloid-β aggregation.

The kinase enrichment results demonstrate the complexity of proteome-wide changes that occur with disease. Among phosphorylation sites with upregulated amyloid-β and tau, the putative sites for nine kinases, three of which are cyclin-dependent kinases including CDK5 which directly phosphorylates tau contributing to the development of AD.(52) Six kinases were under-enriched for activity for amyloid-β and tau including four members of the activin receptor family critical for TGF-β signaling. While TGF-β appears to exacerbate the development of AD, some protective anti-inflammatory effects have also been noted.(53) Similarly, the under-enrichment of members of the GRK family contributes to the complex role of these kinases in AD.(54)

Beyond these concordant effects, the inverted enrichment among multiple kinases between amyloid-β and the other outcomes including tau is fascinating. Protein kinase C is a great example, which showed an enrichment for kinase activity in relation to amyloid-β load, but an under-enrichment for activity in relation to tau tangle burden. Protein kinase C is known to drive an increase in the release of soluble APP along the non-amyloidogenic pathway reducing amyloid-β load.(55,56) In contrast, protein kinase C also directly phosphorylates tau and could actually drive tangle pathology,(57,58) although the direction of these effects has been debated.(59) It appears that protein kinase C regulates GSK3 in a way that reduces some phosphorylation events at S202 and T205, but also enhances phosphorylation at Thr231,(58) a site with particular disease relevance, while independently driving phosphorylation of tau at serine 313 in a manner that contributes to dysfunction.(60) Our results suggest that the counter-balance between these key neuropathologies presents an important challenge when therapeutically targeting protein kinase C (or the other kinases showing this pattern) because therapeutic manipulation may drive counteracting effects on amyloid-β and tau that negate any potential benefits. Further mechanistic work is warranted to better understand these competing pathways in the context of AD pathogenesis.

This study has numerous strengths. To our knowledge, this is the first proteome-wide study on the association between multiple PTMs in the brain and AD neuropathology and cognitive performance. We were able to analyze the effects of PTMs in a well-characterized cohort that includes decades of longitudinal cognitive data prior to death. Despite these strengths, this study was limited by being from a unique subset of non-Hispanic whites (relatively old, impaired, and highly educated), limiting the generalizability to the general non-Hispanic white population, not to say anything of people from other ancestral backgrounds. Further, while this study did characterize four types of PTMs and their relation to AD, there are others that are implicated in AD pathogenesis but were beyond the scope of the present study.

Additionally, to avoid confounding of the PTM relative abundance profiles with their parent protein profile, the applying the adjustment procedure by regressing out protein abundances from the PTM abundances. However, some proteins have different behaviors or different regions. An obvious case is the APP protein where the core of the protein behaves differently from amyloid-β proteolytic fragment that undergoes aggregation and accumulation. Besides the proteolytic processing the situation can get complicated by the presence of the slice isoforms. Therefore, this analysis represents just a first step in the discovery process. In each specific case of the significant PTM it requires follow-up analysis and studies to understand the nature and the mechanism behind the association.

In conclusion, the results from this study indicate that widespread changes to the proteome are observed in the AD brain and relate to both the neuropathology and clinical manifestation of disease. We present novel sites on APP and MAPT as therapeutic targets, as well as novel proteins not previously implicated in AD that have potential to serve as targets for future intervention strategies. Additionally, we explored what kinases may be driving the phosphorylation results, further demonstrating the complex proteome changes in the AD brain. Future studies should focus on diversifying the populations represented by the samples, expanding the scope of PTMs analyzed, and mechanistically validating the targets identified in our study.

## Acknowledgements

Rush Memory and Aging Project is supported by National Institute on Aging grants P30AG10161, P30AG72975, R01AG17917, R01AG015819, U01AG072572, and U01AG046152. Additional support includes National Institute on Aging grants R01-AG059716, R01-AG061518, the Vanderbilt Memory and Alzheimer’s Center, and the Vanderbilt Alzheimer’s Disease Research Center (P30-AG086403). ROSMAP resources can be requested at www.radc.rush.edu.

